# Stationary phase KCl levels trigger changes in Dps conformation to facilitate its partitioning between reversibly-aggregated deposits and Dps-DNA condensates

**DOI:** 10.64898/2026.08.21.746236

**Authors:** Mehak Mahajan, Archit Gupta, Purnananda Guptasarma

## Abstract

In *E. coli* populations subjected to starvation, the transition of cells into stationary-phase is characterized by reduced KCl levels, cytoplasmic acidification, and increased production of Dps, the DNA binding protein from starved cells. Here, we show that at KCl concentrations peculiar to the stationary-phase, Dps displays greater surface hydrophobicity, lower helical content, lower stability to denaturation, greater susceptibility to proteolysis, and an intriguing ability to undergo ‘reversible’ deposition into liquid-solid phase separated (LSPS) aggregates, when no DNA is present. Whereas we have already previously shown that, at growth-phase KCl concentrations, Dps either remains soluble or forms liquid-liquid phase separated (LLPS) condensates with DNA, when DNA is present, here we show that the coacervation of Dps with DNA is exacerbated by stationary-phase KCl concentrations. This leads us to suggest the following interesting possibilities: (i) newly-produced Dps is reversibly kinetically-partitioned between LSPS and LLPS states, at low KCl concentrations, depending on the availability of DNA; (ii) excess Dps that is not coacervated with DNA in the LSPS state, exists in the LSPS state at low KCl concentrations; and (iii) Dps in the LSPS state dissolves to become instantaneously available for the packaging of newly-produced DNA, once nutrition becomes available, growth resumes and cytoplasmic KCl concentrations rise, before there is any production of the growth-phase DNA-packaging protein, HU.

**Importance:** We demonstrate conformational behaviour with physiological implications involving Dps (DNA binding protein from starved cells). We have earlier shown that the highly-soluble Dps forms liquid-liquid phase separated (LLPS) condensates with DNA, at KCl concentrations peculiar to growth-phase *E. coli*. Here, we show that at KCl concentrations peculiar to stationary-phase *E. coli*, Dps undergoes conformational changes to either display heightened LLPS with DNA, or form liquid-solid phase separated (LSPS) aggregates. LSPS aggregates readily dissolve when KCl concentrations are restored to levels peculiar to growth-phase *E. coli*. We propose, therefore, that a dynamic kinetic partitioning of Dps between LSPS and LLPS states maintains excess (i.e., non-DNA-bound) Dps in the form of deposits that dissolve to support packaging of all newly-produced DNA while cells transition back into growth phase (and KCl levels rise), and while the nucleoid-associated protein (NAP) specific to the growth phase, known as HU, is not yet available.

## Introduction

With the decreasing availability of nutrients in the environment, *E. coli* cultures transition from the growth phase to the stationary phase (1). Several changes occur during this transition. Intracellular KCl concentrations drop from ∼250 mM (or higher) to 50 mM (or lower), accompanying changes in intracellular concentrations of NaCl (typically ≤ 5 mM) that rise to match extracellular NaCl concentrations. Concomitantly, intracellular pH drops from ∼ 7.0 to ≥ 5.0 (2). In addition, the intracellular abundance of a DNA-binding, nucleoid-associated protein (NAP) called Dps (or the ‘DNA binding protein from starved cells’), specific to the stationary phase, increases from ∼ 6000 monomers/cell to over 180,000 monomers/cell (3, 4). This rise is the level of Dps parallels a drop in abundance of an NAP called HU (a ‘histone-like protein’), specific to the logarithmic growth phase. As cells transition to stationary phase, HU levels reduce, even as Dps levels rise, allowing Dps to replace HU upon the *E. coli* genome (3). DNA binding by Dps has been reported to increase with reducing KCl concentration, based upon observations made using 50 mM and 150 mM KCl solutions, at pH 7.5 (5). A reduction of pH to 5.0 has also been reported to increase association of Dps with DNA (2, 6, 7). It would appear, therefore, that the drop in intracellular KCl concentration during the transition to stationary phase, and the drop in pH, promote the association of the freshly-synthesized Dps with genomic (and other) DNA.

Recently, we have shown that HU and Dps, both individually and in partnership, engage in mutually-compatible liquid-liquid phase separation (LLPS) behaviour with DNA, which is triggered by mutual macromolecular crowding, and mutual charge neutralization, occurring between the NAP (i.e., HU, or Dps) and DNA (4). Super-resolution microscopic evidence for LLPS-like behaviour in bacterial nucleoids (visualized using overexpressed RFP-HU-A fusion protein), and visual evidence demonstrating the compaction of DNA upon addition of NAPs, demonstrates that LLPS serves as a generic mechanism for the compaction of DNA into nucleoids, and for the maintenance of this compacted state(s) through replication, transcription and cell division, across bacterial cell generations (4).

Given the above, the work presented here was performed to obtain insights into how Dps itself behaves under physico-chemical conditions extant during stationary phase. Studies were performed to examine the conformation, and stability, of Dps as a function of reducing KCl concentration (or pH), in the absence, and presence, of DNA, with a view to understanding whether Dps conformation is altered, and also whether LLPS-mediated coacervation of Dps and DNA is influenced by the lowering of KCl concentration. Notably, a lower pH could exert effects upon Dps, in principle, through changes in net surface charge affecting the protein’s association with DNA (5,7). We report that only a minor conformational change occurs in Dps upon lowering of pH from 7.4 to 5.0. In contrast, profound changes occur upon lowering of KCl concentration (especially down to ∼20 mM). Our results indicate that there is a ‘flaring out’ of polypeptide chain regions on the surface of dodecameric/trimeric Dps, associated with reduced helical content, exposed surface hydrophobicity, increased proteolytic susceptibility and a tendency to undergo reversible agglomeration and precipitation that is associated with a reversible increase in hydrodynamic radius, suggestive of reversible associations amongst partially-unfolded multimers of Dps.

Such behaviour could involve either the association of dodecamers, or of trimers derived through dissociation facilitated by lowering of KCl concentration; similar to trimers of Dps already reported in a different context (8). We also report that the ability of Dps to undergo LLPS with DNA increases with reducing KCl concentration. During lowering of KCl concentrations from 300 mM to 20 mM, the interesting differences between (i) the ‘reversible precipitation’ of Dps, seen at high protein concentrations, in the absence of DNA, at KCl concentration ≤ 50 mM, and (ii) the increased ‘LLPS’ of Dps with DNA, seen at high protein concentrations, in the presence of DNA, suggests that excess Dps probably deposits reversibly into DNA-lacking aggregates and precipitates that serve as a sink from which Dps is later sourced, once stationary phase ends, and KCl concentrations rise.

## Results and Discussion

### When KCl drops to 20 mM, at low protein concentrations, Dps shows altered diameter, hydrophobicity, conformation, and stability

#### Delayed SEC elution at 20 mM KCl

**Figure 1A** shows that Dps elution from a Superdex 200 Increase size exclusion chromatography (SEC) column shifts to a larger elution volume, when KCl concentration is decreased from 150 mM KCl to 20 mM KCl. This indicates two possibilities: (1) dissociation of Dps dodecamers into smaller species; and/or (2) partial unfolding of Dps chains with, or without, accompanying dissociation, causing exposure of surfaces that interact hydrophobically with the column, to delay elution. Notably, delayed elution due to hydrophobic protein-resin interactions have previously been reported (9–11).

**Figure 1.**
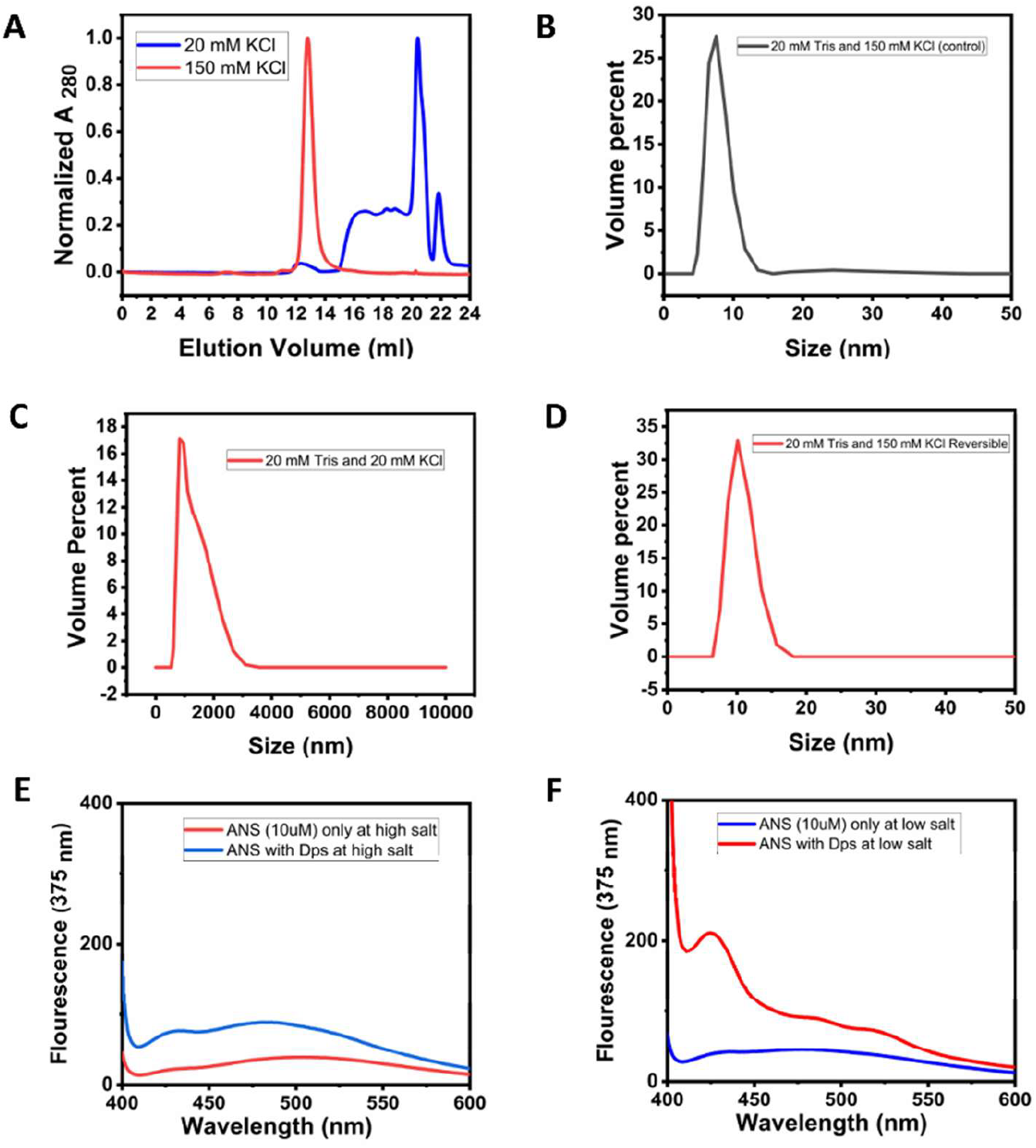
Assessment of Dps size and hydrophobicity at 150 mM and 20 mM KCl. *A*, Differences in elution volume of Dps on a Superdex 200 Increase column in 150 and 20 mM KCl. *B*, DLS-based measurement of Dps hydrodynamic diameter in 150 mM KCl. *C*, DLS-based measurement of Dps hydrodynamic diameter in 20 mM KCl. *D*, DLS-based measurement of Dps hydrodynamic diameter upon restoration of KCl concentration to 150 mM. *E*, Fluorescence spectra of ANS in control samples lacking Dps (red) and in the presence of 0.5 mg/mL Dps, in 150 mM KCl (blue). *F*, Fluorescence spectra of the hydrophobic reporter dye, ANS, in control samples lacking Dps (blue) and in the presence of 0.5 mg/mL Dps, in 20 mM KCl.

#### A reversible 100-fold increase of hydrodynamic diameter at 20 mM KCl

A differential diagnosis between ‘dissociation’ and ‘partial unfolding’ can be facilitated by dynamic light scattering (DLS). **Figures 1B, and 1C**, respectively, show that the hydrodynamic diameter of Dps changes from ≤10 nm at 150 mM KCl, to ≥1000 nm at 20 mM KCl, indicating that dissociation into smaller species is not responsible for the delayed elution. Interestingly, as seen in **Figure 1D**, the increase of hydrodynamic diameter to ≥1000 nm, at 20 mM KCl, is entirely reversed through restoration of KCl concentration to 150 mM.

#### Enhanced surface hydrophobicity at 20 mM KCl

The hydrophobic dye, 8-anilino-1-naphthalene sulfonate (ANS), is a well-known reporter of hydrophobic surfaces in proteins since ANS binds to such surfaces through its naphthalene moiety to display enhanced fluorescence intensity and/or blue-shifting of emission (12). **Figures 1E, and 1F**, respectively, show that while ANS shows no sign of binding to Dps at 150 mM KCl, there is both an enhancement and a blue-shifting (to ∼425 nm) of the dye’s fluorescence emission, at 20 mM KCl, indicating the exposure of hydrophobic surfaces upon Dps due to the drop to 20 mM KCl.

#### The exposed surface hydrophobicity leads to agglomeration and protein-resin interactions

Together, the SEC and DLS data presented above suggest that the lowering of KCl from 150 mM to 20 mM, causes Dps to (i) display larger average hydrodynamic volume(s), potentially through partial unfolding of chain sections and/or formation of soluble agglomerates/aggregates, and (ii) expose surface hydrophobicity, that promotes chain-chain associations and also leads to binding protein-resin interactions and delayed SEC elution. Next, we used low-resolution spectroscopic methods to examine Dps behaviour at 150 mM and 20 mM KCl.

#### Reduced overall helical content, at 20 mM KCl

To examine whether conformational changes occur in Dps between 150 mM and 20 mM KCl, we performed circular dichroism (CD) spectroscopy. **Figure 2A** shows a progressive loss of alpha helical structural content as a function of reducing KCl concentration, evident from lowering in intensity of the mean-residue ellipticity (MRE) signal between 250 and 200 nm (and at the 222, and 208, nm band positions ascribable to alpha helical structural content). Comparisons of normalized spectral curves also indicate changes in the shape of spectra (**Supplementary Figure S1**).

**Figure 2.**
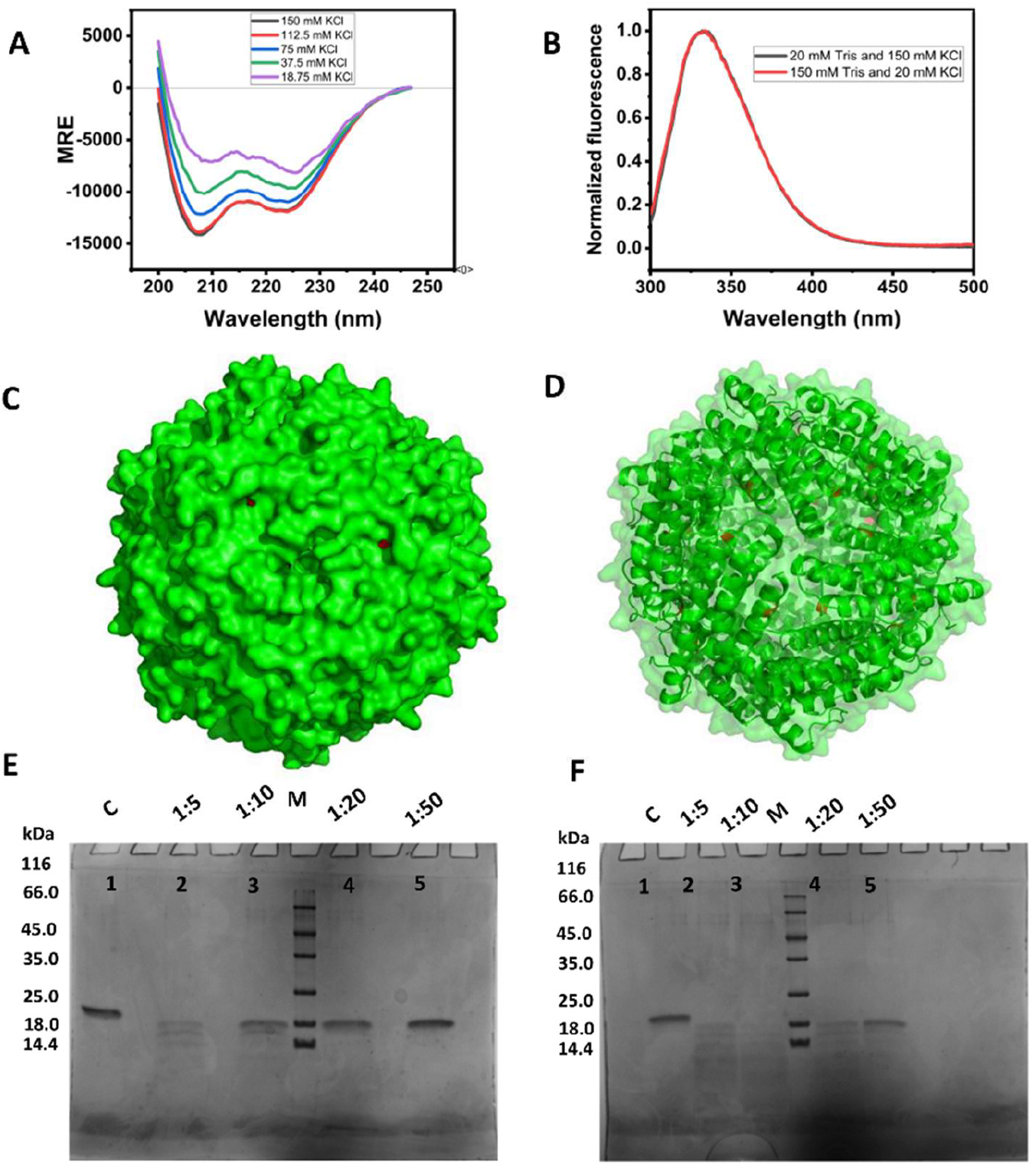
Assessment of Dps structure and proteolytic susceptibility at 150 mM and 20 mM KCl. *A*, CD spectra of Dps in decreasing concentrations of KCl (mentioned in the inset). *B*, Fluorescence spectra of Dps in 20 mM (black) and 150 mM KCl (red). *C*. All-atom representation of the Dps dodecamer’s surface, with atoms belonging to tryptophan residues (in red) largely invisible on account of being buried. *D*. All-atom representation of the Dps dodecamer shown with 60% transparency, and superimposition of a ribbon-diagram representation of the backbones of monomers, providing a visual assessment of the burial of tryptophan residues (in red) inside the dodecamer’s core. *E*. Subtilisin A based assessment of the proteolytic susceptibility of Dps in 150 mM KCl. *F*. Subtilisin A based assessment of the proteolytic susceptibility of Dps in 20 mM KCl. In both panels *E* and *F*, individual lanes are annotated in respect of the molar ratio of subtilisin:Dps used for the reaction.

#### No change in tryptophan environments, at 20 mM KCl

The wavelength of maximal protein fluorescence emission (^em^λ_max_) responds acutely to changes in tryptophan residue environment polarity. Dps has two tryptophans, W52 and W160. We examined whether the changes in Dps conformation indicated by changes in size, hydrophobicity and helical content are also reflected in changes in ^em^λ_max_. **Figure 2B** shows that Dps fluorescence at 150 mM and 20 mM KCl is identical, and superimposable. This establishes that the environments of W52, and W160, remain unperturbed. This result is compatible with the observed lack of dissociation at 20 mM KCl. Any profound loss of structure associated with dissociation would be expected to have altered the ^em^λ_max_. This result is also compatible with the flaring out of outer regions of Dps in the multimeric state, to expose hydrophobicity and facilitate agglomeration/aggregation, because this could be envisaged to leave the structure around W52 and W160 unperturbed, at 20 mM KCl. **Figures 2C, and 2D**, respectively, show ribbon diagram views of the Dps dodecamer (in green) with 0 % and 60 % transparency settings. Tryptophan residues (in red) are visible in both views, from which it is clear that the 24 tryptophan residues of Dps (2 per chain; 12 chains) are buried within the dodecamer’s core, facilitating conception of conformational changes occurring in the outer helices without affecting the molecule’s core, at low KCl concentrations.

#### Greater proteolytic susceptibility, at 20 mM KCl

In **Figures 2E, and 2F**, respectively, we show results from an experiment involving SDS-PAGE-based assessment of Dps degradation by the non-specifically-acting protease, subtilisin (3.4 µM dodecamer, or 0.8 mg/mL Dps; using protease:Dps molar ratios of 1:50, 1:20, 1:10 and 1:5, over 120 minutes, in 50 mM, and 20 mM, KCl). The data, in terms of the intensities of surviving bands under different conditions, establishes that Dps survives proteolysis to a far greater degree at 150 mM KCl, than at 20 mM KCl. The distinction is most evident in gel lanes corresponding to a protease:Dps ratio of 1:10. At a protease:Dps ratio of 1:5, no residual Dps is seen at either KCl concentration; however, even here, a greater number of bands corresponding to partially-degraded Dps is seen at 20 mM KCl, compared to 150 mM KCl, indicating higher proteolytic susceptibility at 20 mM KCl, presumably owing to some (partial) unfolding of outer helices.

#### Less thermal stability, at 20 mM KCl

The sensitivity of the ^em^λ_max_ of a protein to the polarity of the environments of tryptophan residues causes the ^em^λ_max_ to shift to longer wavelengths as any protein population unfolds, causing the ^em^λ_max_ to shift to 352-353 nm for completely unfolded protein, with accompanying changes in the ratio of fluorescence intensities at 330 nm and 350 nm, which can be plotted to monitor unfolding. **Supplementary Figures S2A, and S2B**, respectively, show plots of Dps 330/350 nm fluorescence ratios in 150 mM, and 20 mM KCl, as a function of temperature (raised to elicit unfolding). The data shows that Dps is significantly less thermally-stable at 20 mM KCl than at 150 mM KCl, with changes seen in the 330/360 nm ratio at lower temperatures in the 20 mM KCl plot.

#### Gdm.HCl initially restores structural content, before unfolding Dps

**Supplementary Figures S2C, and S2D**, respectively, show that at 20 mM KCl, structural content increases with increasing Gdm.HCl concentration up to 1 M. This is presumably because of the increase in ionic strength, due to increase in Gdm.HCl concentration acting as a proxy for increase in KCl concentration. Of course, structural content then decreases upon further increase in Gdm.HCl concentration, due to chain unfolding. With the 150 mM KCl sample, the same phenomenon is seen, except that the structural content (which is higher to begin with) shows signs of decreasing at higher Gdm.HCl concentrations, as expected for a more stable protein.

### When KCl drops to 20 mM, at high protein concentrations, Dps shows precipitation which is reversed by raising KCl to 150 mM

#### An uncommon, reversible agglomeration of Dps through partial unfolding

The studies presented in the above section used low to moderate Dps concentrations (0.2 to 0.5 mg/mL, i.e., well below the concentrations of ∼5.6 to 9.3 mg/mL found in stationary-phase *E. coli*). Since we saw evidence of reversibility in soluble Dps agglomerates/aggregates at low Dps concentrations, we decide to examine whether (a) an increase in protein concentration transforms soluble Dps agglomerates/aggregates into insoluble aggregates/precipitates, and also whether (b) precipitates (assuming that they are formed) are solubilized back into dodecameric Dps, through restoration of KCl to 150 mM. Notably, any observed reversibility would need to be explained by variations in surface charge, powerful enough to overwhelm hydrophobic interactions causing the aggregation and precipitation. Thus, we performed multiple types of experiments to investigate this, as described below.

#### With increasing protein concentration, progressive agglomeration/aggregation is seen only at 20 mM KCl (and not at 150 mM KCl)

We varied Dps protein concentration from 0.2 mg/mL to 1.6 mg/mL, monitoring solution turbidity (optical density; O.D._600_) as a reporter of agglomeration, aggregation, condensation, or precipitation. **Figure 3A** shows that turbidity increases steadily with Dps concentration, at 20 mM KCl, but not at 150 mM KCl. This establishes that chain associations are promoted by decreasing KCl concentrations.

**Figure 3.**
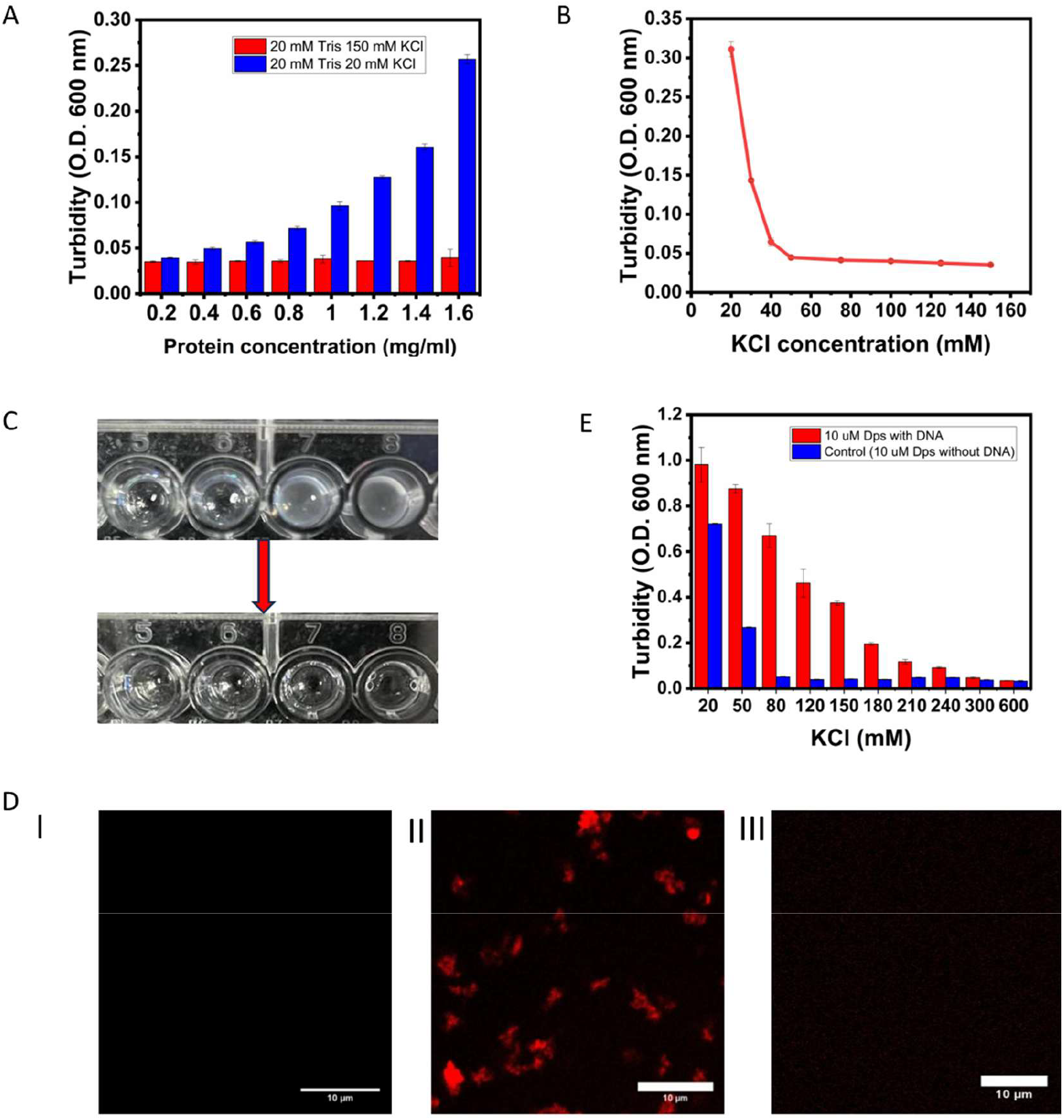
Assessment of Dps agglomeration/aggregation/precipitation in DNA’s absence, and liquid-liquid phase separation (LLPS) in the presence of DNA. *A*, Turbidity (O.D. 600) as a function of increasing Dps concentration, in 150 mM KCl (shown in red) and 20 mM KCl (shown in blue). *B*, Turbidity (O.D. 600) of 2.4 mg/mL Dps samples as a function of increasing KCl concentration. *C*, Turbidity in 50, 40, 30 and 20 mM KCl samples (left-to-right; 4 wells; top panel) disappearing upon restoration of KCl concentration to ∼150 mM in all wells (bottom panel). *D*, Confocal microscopic images of 2.4 mg/mL Dps in 150 mM KCl (left panel), 20 mM KCl (middle panel), and upon restoration of KCl concentration to 150 mM (right panel), revealing aggregates/precipitates only in the 20 mM KCl sample of Dps. *E*, Turbidity (O.D. 600) of 2.4 mg/mL (∼10 µM) Dps samples as a function of increasing KCl concentration, in the presence of 0.15 mg/mL salmon testis DNA (shown in red) and absence of DNA (shown in blue).

#### With varying KCl concentration, at high protein concentration, progressive agglomeration/aggregation is seen as KCl is lowered from 150 mM to 20 mM

We varied KCl concentration from 150 mM to 20 mM, using 2.4 mg/mL Dps (10 µM; a higher protein concentration, but still a fraction of the stationary-phase Dps concentration of ∼ 5.6 – 9.3 mg/mL). **Figure 3B** shows that turbidity increases steadily as a function of decreasing KCl concentration, with significant turbidity seen only between 50 and 20 mM KCl.

#### Turbidity at high protein concentration in 20 mM KCl is reversed by raising KCl to 150 mM

We took turbid solutions formed by 2.4 mg/mL Dps, at KCl concentrations of 50, 40, 30, and 20 mM, and then imaged these solutions (each in a volume of 100 µl) in separate wells of a 96 well-plate. We then increased the KCl concentration to 150 mM KCl in well, by adding 5.40, 5.94, 6.48 and 7.02 µl KCl, respectively (from a 2.0 M stock), to raise KCl concentration to 150 mM with negligible dilution of protein concentration. **Figure 3C** shows that there is an disappearance of turbidity in all wells.

#### Insoluble precipitates formed at high protein concentration, in 20 mM KCl, are also solubilized by raising KCl to 150 mM

We used fluorescence microscopy to examine, and image, 2.4 mg/mL Dps at 150 mM KCl and 20 mM KCl, before, and after, conversion of the 20 mM solution to a 150 mM solution. **Figure 3D** shows that (i) there are no precipitates at 150 mM KCl; (ii) there is formation of precipitates at 20 mM KCl; and (iii) formed precipitates are solubilized through restoration to 150 mM KCl.

### In 20 mM KCl, at high protein concentrations, Dps and DNA undergo enhanced liquid-liquid phase separation

#### While Dps aggregates/precipitates create turbidity only between 50 mM and 20 mM KCl, Dps condensates (LLPS) cause turbidity between 300 mM and 20 mM KCl

As mentioned earlier, we have already previously shown that Dps and DNA undergo mutual liquid-liquid phase separation (LLPS) under conditions simulating macromolecular crowding inside bacteria (4). **Figure 3E** shows that when 4 % w/v PEG is present, the excluded volume effect mildly increases the agglomeration/aggregation of Dps at low ionic strength. For a 2.4 mg/mL Dps solution, although there is only a negligible difference in turbidity between 80 and 50 mM KCl in the absence of PEG, as seen in **Figure 3B**, the presence of PEG significantly increases difference in turbidity between 80 mM and 50 mM KCl, as seen in **Figure 3E**. This is because, with DNA present in solution, the level of formation of Dps-DNA condensates increases as KCl concentration is reduced from 600 mM to 20 mM, due to the occurrence of LLPS between Dps and DNA (here sourced from salmon testis). When DNA is present, phase separation can be seen to dominate development of turbidity over the entire range of KCl concentrations, including at 50 mM and 20 mM KCl (for which agglomeration/aggregation, but not LLPS, is seen in the absence of DNA). **Supplementary Figure S3** shows that when DNA is present, at 20 mM KCl, turbidity derives from the formation of largely spherical LLPS droplets. However, when DNA is absent, turbidity derives from the formation of irregularly-shaped Dps agglomerates/aggregates and precipitates.

All of the above data was collected at pH 7.4. Incidentally, we also examined the effect of independently lowering pH from 7.4 to 5.0, without any concomitant lowering of KCl concentration (i.e., by maintaining KCl at 150 mM). **Supplementary Figure S4** presents the CD spectroscopic data for this study. The effect of lowering of pH upon Dps structural content was a slight increase in structural content. While we did not follow this up with detailed experiments at pH 5.0 (since, due to the use of a different buffer, there would be a need to examine ionic strength changes in more nuanced fashion), we report that at first glance it seems that the reduction in pH marginally increases structural content in Dps; however, to a much lower extent than structure is lost with reduction in KCl concentration, especially below ∼50 mM KCl. The question of whether the drop in pH manages to somewhat counter the effects of a drop in KCl concentration needs to be followed up in more detail.

## Conclusions

We show that at KCl concentrations below 50 mM, and at a relatively-low Dps concentration of 0.2 mg/mL (below physiological Dps concentrations in stationary-phase cells), Dps undergoes partial structural destabilization, associated with lower helical content, increase in hydrodynamic radius, exposure of hydrophobicity, chain associations that are reversible, and greater proteolytic susceptibility. We also show that when Dps concentration is increased to between ½ and ¼ physiological Dps concentration, there is a progressive (but reversible) increase in turbidity associated with precipitation, at Dps concentrations between 0.4 and 2.4 mg/mL.

Thus, at all protein concentrations, increased turbidity due to agglomeration, aggregation, or precipitation, is seen when KCl is ≤50 mM. However, when DNA is present, phase-separation is seen at all KCl concentrations, with progressive increase in turbidity due to phase-separation as KCl concentration is lowered from ∼300 mM to 20 mM KCl. The turbidity due to LLPS at 20 mM KCl in the presence of DNA is ∼2.5 times greater than that seen at 150 mM KCl. This turbidity is also over ∼1.4 times greater than that owing to Dps aggregation and precipitation at 20 mM, in the absence of DNA.

Our results raise the possibility that conformational changes in Dps facilitate improved phase-separation at low KCl concentrations and cause reversible precipitation of any excess Dps into deposits that act as ‘sinks’, when enough DNA is not present for coacervation with Dps. Such Dps ‘sinks’ could do more than just remove excess protein from solution. Since they redissolve with rise in KCl concentration, they could act as an immediate source of Dps for coacervation with newly synthesized DNA, prior to fresh synthesis of HU, upon resumption of growth, during early stages of restoration of stationary-phase populations back to logarithmically-growing cells (upon release of starvation). Such behaviour would parallel the concentration-buffering role proposed for LLPS, in which a dense phase forming above a saturation concentration holds the surrounding free concentration constant (13, 14). In other words, the saturation thermodynamics that applies to LLPS coacervates could also apply to DNA-lacking Dps sinks, such that Dps synthesized beyond the capacity for DNA binding could potentially be ‘sunk’ into reversible deposits, without over-compacting the genome, and keeping DNA-binding cellular Dps levels constant for any given level of intracellular KCl, during the descent into stationary-phase. With rise of KCl levels accompanying resumption of growth triggered by availability of nutrition, Dps stored in such precipitates would obviously be released immediately, and used. The large amount of Dps synthesized during stationary phase could thus function as an adjustable reserve to calibrate genome compaction with cellular KCl levels and nutritional status.

## Materials and Methods

### Dps expression, purification and stocks

C-terminal 6xHis affinity tag-displaying Dps was produced, expressed and purified, as described previously, in *Escherichia coli* strain BL21 (DE3) pLysS* (4). Yields from 1 L cultures (7 to 9 mg Dps) saturated the Ni-NTA resin (1.5 ml) used for affinity purification, removing need for further purification. Imidazole was removed through buffer replacement involving repeated additions of 20 mM Tris, 150 mM KCl, buffer, and centrifugal membrane-based concentration (10 kDa cutoff) at 4500 rpm. Aliquots were frozen and cryopreserved at – 80 °C.

### DPS concentration estimation

Dps absorbance was measured at 280 nm, with an O.D_280_ of 0.83 corresponding to 1 mg/mL Dps (for an Є of 15470 M^-1^ cm^-1^). A solution of 1 mg/mL contains 4.5 µM dodecameric Dps, and 54.5 µM monomeric Dps.

### ANS assay for hydrophobicity

The hydrophobic dye, 8-Anilino-1-naphthalenesulfonic acid (ANS), was used to probe exposed surface hydrophobicity, by monitoring ANS fluorescence which increases and displays blue shifts upon binding to hydrophobic protein patches, upon 375 nm excitation (emission measured between 400 and 600 nm). ANS (working concentration 1 uM) was incubated with Dps (0.5 mg/mL) in 100 µl reactions, in 150 mM, or 20 mM KCl.

### Circular Dichroism (CD)

Structural content in Dps was examined using a MOS-500 CD spectrometer (BioLogic, France). Dps (0.4 mg/mL) was incubated in 20 mM Tris, pH 7.4, containing varying concentrations of KCl. Raw ellipticity was converted to mean residue ellipticity (MRE), using standard formulae. Spectral normalization was carried out using MRE values at 216 nm, to examine shape changes.

### Fluorescence spectroscopy

Fluorescence emission was used to examine Dps (0.5 mg/mL) in 20, and 150 mM KCl, using excitation at 295 nm wavelength (emission measured between 300 and 500 nm), using a 0.3 x 0.3 cm cuvette, and a Varian Cary Eclipse fluorimeter (Agilent Technologies).

### Size exclusion chromatography

Dps protein (500 µl; 0.2 mg/mL) in 20 mM Tris, pH 7.4 containing 20 mM KCl, or 150 mM KCl, was loaded on a GE Superdex-200 Increase size exclusion chromatographic column connected to a GE AKTA Purifier-10 workstation. For chromatography at different KCl concentrations, pre-equilibrated was done using buffer of the appropriate KCl concentration.

### Fluorescence labelling

The cysteine mutant of Dps (Dps S22C) was labelled with maleimide dye [Alexa Fluor 594 C^5^ Maleimide], through incubation of 88 µM Dps with 168 µM TCEP (Tris(2-carboxyethyl) phosphine hydrochloride; Merck) and 168 µM dye, at 37 °C for 3 hours. Free dye was removed through centrifugal ultrafiltration. Labelling efficiency was calculated using standard procedures.

### Dynamic light scattering (DLS)

Dps (0.3 mg/mL; 100 µl) in 20 mM, and 150 mM KCl, was analysed on a Zetasizer Nano ZS90 (Malvern Instruments, Malvern, UK).

### Aggregation Assays

O.D._600_ was used as proxy for aggregation. Assays assessed effects of changing KCl and protein concentration. Dps protein (2.4 mg/mL; 150 µl) in 20 mM Tris, pH 7.4, in the presence of different KCl concentrations varying between 20 and 600 mM KCl, was subjected to O.D._600_ measurements, using a 96-well plate upon a POLARstar Omega plate reader (BMG Labtech, Germany), to assess aggregation. Similarly, Dps (0.2 mg/mL to 1.6 mg/mL, using increments of 0.2 mg/mL; 150 µl volume) was subjected to O.D._600_ measurements in the presence of 20 mM, or 150 mM, KCl. To examine reversibility of aggregation, Dps protein (2.4 mg/mL; 100 µl), in the presence of different concentrations of KCl (20, 30, 40, and 50 mM) was subjected to O.D._600_ measurements, with wells containing aggregates restored to 150 mM KCl through addition of appropriate buffer volumes containing 2M KCl.

### Fluorescence microscopy

To examine Dps aggregation/precipitation in 150 mM or 20 mM KCl, and reversibility of precipitation upon restoration to 150 mM KCl, microscopy was carried out on a Zeiss LSM 980 microscope. Conditions used for assessing LLPS are separately mentioned in the supplementary file, with data.

### Subtilisin sensitivity assay

A working concentration of 3.4 µM Dps was incubated with subtilisin at 4 different ratios (1:5, 1:10, 1:20, and 1:50) of subtilisin:Dps, at 55 °C, for 2 hours, using 20 µl reactions. SDS PAGE was used to examine digestion.

### Chemical unfolding using CD

To unfold Dps using guanidium hydrochloride (Gdm.HCl), 300 µl samples of Dps (0.2 mg/mL) were placed in different concentrations of Gdm.HCl.

## Data availability statement

The Article and Supplementary Information contain all the data for this manuscript.

## Ethics statement

The work presented in this paper uses only commercially sourced reagents, biophysical and biochemical instrumentation, and recombinant protein molecules that have been produced in bacteria by our laboratory. It does not violate any ethical guidelines of any description.

## Supporting information

This article contains supporting information.

## Author contributions

M.M., A.G., and P.G. conceptualization; M.M., A.G., and P.G. validation; M.M., A.G., and P.G. investigation; M.M., A.G., and P.G. writing - original draft; P.G. supervision; M.M., A.G., and P.G. writing - review and editing.

## Acknowledgements: Funding and additional information

We thank the Ministry of Education (Centre of Excellence Grant in Protein Science, Design and Engineering to P.G), the Department of Biotechnology (HEHRC grant to P.G), and TATA Sons (TATA Transformation Prize to P.G.). M.M., and A.G., thank DBT (India), and CSIR (India), respectively, for research fellowships. We thank the Government of India for financial support (for equipment/consumables) extended through IISER Mohali. We thank Soumyajit Kar, Aakanksha Meena for occasional help with experiments.

